# Light-based tuning of ligand half-life supports kinetic proofreading model of T cell activation

**DOI:** 10.1101/432864

**Authors:** Doug Tischer, Orion D. Weiner

**Affiliations:** Cardiovascular Research Institute and Department of Biochemistry and Biophysics, University of California San Francisco, San Francisco, California, USA

## Abstract

T cells are thought to discriminate stimulatory versus non-stimulatory ligands by converting small changes in ligand binding half-life to large changes in cell activation. Such a kinetic proofreading model has been difficult to test directly, as existing methods of altering ligand binding half-life also change other potentially important biophysical parameters, most notably the stability of the receptor-ligand interaction under load. Here we develop an optogenetic approach to specifically tune the binding half-life of a light-responsive ligand to a chimeric antigen receptor without changing other binding parameters. By simultaneously manipulating binding half-life while controlling for receptor occupancy, we find that signaling is strongly gated by ligand binding half-life. Our results provide direct evidence of kinetic proofreading in ligand discrimination by T cells.

**One Sentence Summary:** Direct control of ligand binding half-life with light shows that lifetime, not occupancy, dominates T cell activation.

## Main Text

The T cell response begins when the T cell antigen receptor (TCR) binds its cognate peptide-major histocompatibility complex (pMHC) on an antigen-presenting cell. Ligand discrimination is essential, as falsely recognizing a self-peptide can lead to autoimmunity, while missing a foreign-peptide allows pathogens to evade detection. Compounding this challenge, self-pMHCs generally outnumber foreign-pMHCs by several orders of magnitude (*1*–*4*). How the TCR recognizes its foreign-pMHC while ignoring the more numerous self-pMHC is a fundamental open question in immunology.

For many cell surface receptors, occupancy drives the strength of signaling. Lower affinity ligands can signal as well as high affinity ligands when their concentration is increased such that an equal number of receptors are bound (*5*). This is unlikely to be the case for activation of the TCR, because low affinity pMHCs are significantly less potent than high affinity pMHCs, even when adjusted for occupancy (*6*). Furthermore, given the large excess of self-pMHC (*1*–*4*) and the relatively small differences in binding affinity (*7*–*9*), it is likely that more TCRs are bound to self-pMHC than foreign-pMHC. For a T cell to detect foreign pMHCs, it is likely that some ligand-intrinsic factor makes bound foreign-pMHCs more stimulatory than bound self-pMHCs.

A major model of T cell ligand discrimination is kinetic proofreading (*10*), which predicts that small differences in ligand binding half-life can be amplified into large differences in signaling. Such a model is attractive because it allows a few receptors bound to long-lived ligands to signal better than many receptors bound to short-lived ligands, potentially explaining why abundant self-pMHCs do not activate T cells, though scarce foreign-pMHCs do. It posits a delay made up of multiple irreversible biochemical steps between ligand binding and downstream signaling. Only ligands that bind persistently signal effectively. These steps only occur while the ligand is bound and quickly reset upon dissociation, preventing two successive short binding events from mimicking one long binding event. This mechanism is based on the kinetic proofreading that gives DNA replication, protein translation (*11*, *12*), and mRNA splicing (*13*) specificity far beyond what would be predicted from equilibrium binding constants alone. Consistent with this hypothesis, kinetic measurements generally show that stimulatory pMHCs have longer binding half-lives, while non-stimulatory pMHCs have shorter binding half-lives (*7*– *9*).

Kinetic proofreading effects need to be strongest when differences in pMHC binding affinities are small, as abundant self-pMHCs will bind more TCRs than scarce foreign-pMHCs. Strong kinetic proofreading ensures that short self-pMHC binding events are unlikely to contribute to downstream signaling. However, recent techniques that measure pMHC binding affinities in more native 2D environments show a larger range of binding affinities than previously measured (*14*), suggesting that there may not be as large an excess of self-pMHC bound to the TCR as previously thought and raising the question on whether ligand half-life or receptor occupancy dominates ligand discrimination.

To probe kinetic proofreading in T cells, the field most commonly relies on altered peptide series to change binding half-lives. The limitation of this approach is that when the binding interface is altered, both the bond’s half-life and stability under tension must change together (**Figure 1A**, left). Differentiating the effects of these two variables is essential because recent evidence has shown that the TCR is mechanosenstive (*15*–*17*) and that foreign-pMHC can form catch bonds with the TCR, growing more stable under load (*18*). Thus, conventional mutational approaches make it difficult to determine whether changes in signaling arise from alterations in binding half-life or reflect changes in the force being applied to the TCR. To specifically probe the role of binding half-life in ligand discrimination, the field needs new techniques to manipulate the receptor-ligand half-life while leaving force transmission and all other aspects of the interaction unchanged. Here we develop an optogenetic approach for T cell activation that addresses this need.

**Figure 1.**
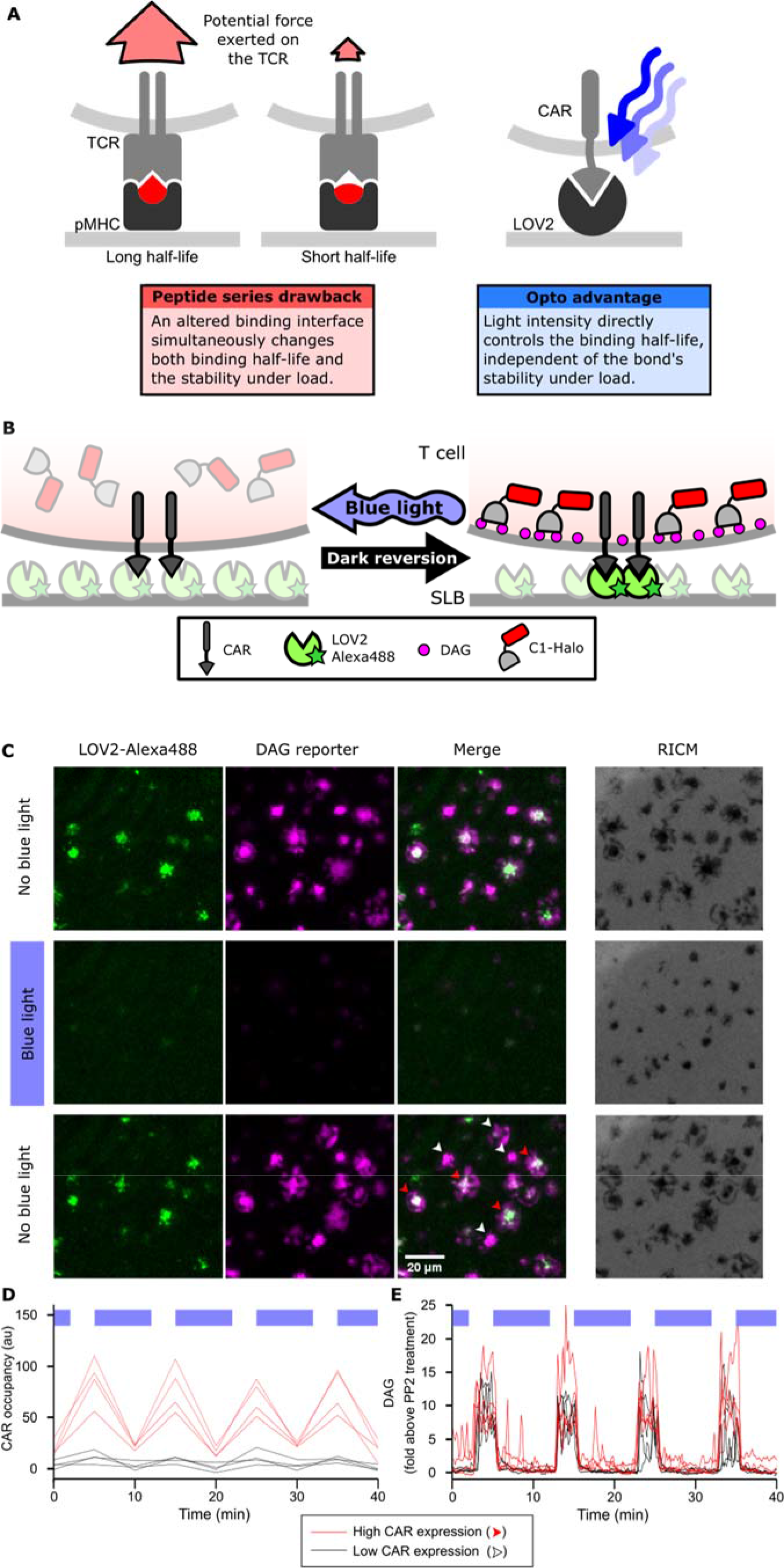
Strategy for testing kinetic proofreading with optogenetic tools. (**A**) Conventional methods of mutating the pMHC to alter the binding half-life also change the binding interface, which changes several parameters at once. By contrast, optogenetic control allows light intensity to control ligand binding half-life while keeping the binding interface constant. Therefore, no other aspects of the receptor-ligand interaction change. Red and blue lines highlight the binding interfaces. (**B**) Schematic of experimental setup. Jurkat cells expressing a live cell DAG reporter and a Zdk-CAR are exposed to an SLB functionalized with purified, dye-labeled LOV2. In the dark, LOV2 binds to and accumulates under the receptor, and stimulates DAG production, recruiting the reporter to the plasma membrane. Blue light excites LOV2, inducing its dissociation from the receptor to terminate signaling. (**C**) Montage from a time course in which cells were alternately stimulated in the presence or absence of blue-light. White arrows highlight cells with low to undetectable receptor occupancy, and red arrows highlight cells with high receptor occupancy. (**D**and **E**) Cells with very different levels of receptor occupancy (**D**), can have similar DAG levels (**E**), suggesting that receptor occupancy is not a good predictor of DAG levels. Top blue bars indicate the presence of blue light.

For optogenetic control of ligand half-life and T cell activation, we constructed a chimeric antigen receptor (CAR) that binds its ligand in a light-dependent fashion. CARs are single chain mimetics of the native, multi-subunit TCR complex and minimally consist of an extracellular binding domain, a transmembrane domain and an intracellular signaling domain. Because CARs are not restricted to binding pMHCs, they can be engineered to signal in response a wide variety of ligands by changing the extracellular domain. Once bound, they initiate downstream signaling by becoming phosphorylated and recruiting Zap70 to the receptor (*19*).

For generating a light-responsive ligand, we choose the LOVTRAP system (*20*) because it is one of only two optogenetic systems where light drives the dissociation of two proteins on the order of seconds, the approximate physiological range of pMHC half-lives (*7*–*9*). By presenting the blue-light sensitive protein LOV2 on a supported lipid bilayer (SLB) and fusing its binding partner (Zdk) to the extracellular domain of a CAR, we generated a CAR with a light-gated ligand. In the dark, the Zdk-CAR binds the LOV2 ligand with nanomolar affinity (*20*). When excited by a photon of blue-light (<520 nm), LOV2 changes conformation and unbinds the receptor. Thus, the intensity of blue light controls the length of time individual molecules of LOV2 and the CAR interact. High intensity blue light enforces a short binding half-life, while low intensity blue light enforces a long binding half-life. This optogenetic approach enables one receptor-ligand pair to explore a range of binding half-lives while maintaining an identical binding interface, ensuring the bond’s stability under tension remains unchanged. (**Figure 1A**, right).

We first validated the ability of the LOV2 ligand to photoreversibly bind the Zdk-CAR. Clonal Jurkat cells stably expressing the Zdk-CAR were exposed to SLBs functionalized with purified Alexa-488-labeled LOV2 (**Figure 1B**). Because LOV2 diffuses freely in the bilayer and becomes trapped upon interaction with the Zdk-CAR, we can measure receptor occupancy by the accumulation of LOV2 under the cell. As expected, LOV2 accumulated under the cells in the absence of blue light and dispersed following illumination with blue light (**Figure 1C**, Videos 1 and 2). Blue light drives multiple cycles of binding and unbinding without apparent loss of potency (**Figure 1D** and **Figure 1–figure supplement 1A**).

We next verified that binding of the LOV2 ligand induces cell signaling through the CAR. We measured accumulation of the lipid diacylglycerol (DAG) by quantifying the translocation of the C1-domains from protein kinase C theta from the cytosol to the plasma membrane with Total Internal Reflection Fluorescence (TIRF) Microscopy (**Figure 1B**). DAG levels spiked and cells spread onto the SLBs in a blue-light-gated fashion (**Figure 1C, E** and Video 3). These data confirm that the photoreversible binding of LOV2 to the Zdk-CAR leads to photoreversible signaling. Low DAG levels in the presence of blue light was not due to cells detaching from the SLB, as cells were passively adhered with an anti-β2 microglobulin antibody (Video 4), and Reflection Interference Contrast Microscopy (RICM) indicated that the cells remained in continuous contact with the SLB (**Figure 1C**, **Figure 1–figure supplement 2** and Video 5).

We noticed that within the same field of view, cells could have high or low-to-undetectable receptor occupancies (red and white arrows, respectively; **Figure 1C, D**). However, despite clear differences in receptor occupancy, all cells had similar DAG levels (**Figure 1E**). This result suggested that something other than receptor occupancy dominates downstream signaling.

As blue light affects the LOV2-Zdk binding half-life, we sought to measure this relation in more detail. The release of purified Zdk from SLBs functionalized with LOV2 was measured upon acute exposure to different blue light intensities (**Figure 2A** and **Figure 2–figure supplement 1**). Increasing blue-light intensities increased Zdk dissociation and tuned the effective binding half-life from ten seconds to approximately 500 ms (**Figure 2B**).

**Figure 2.**
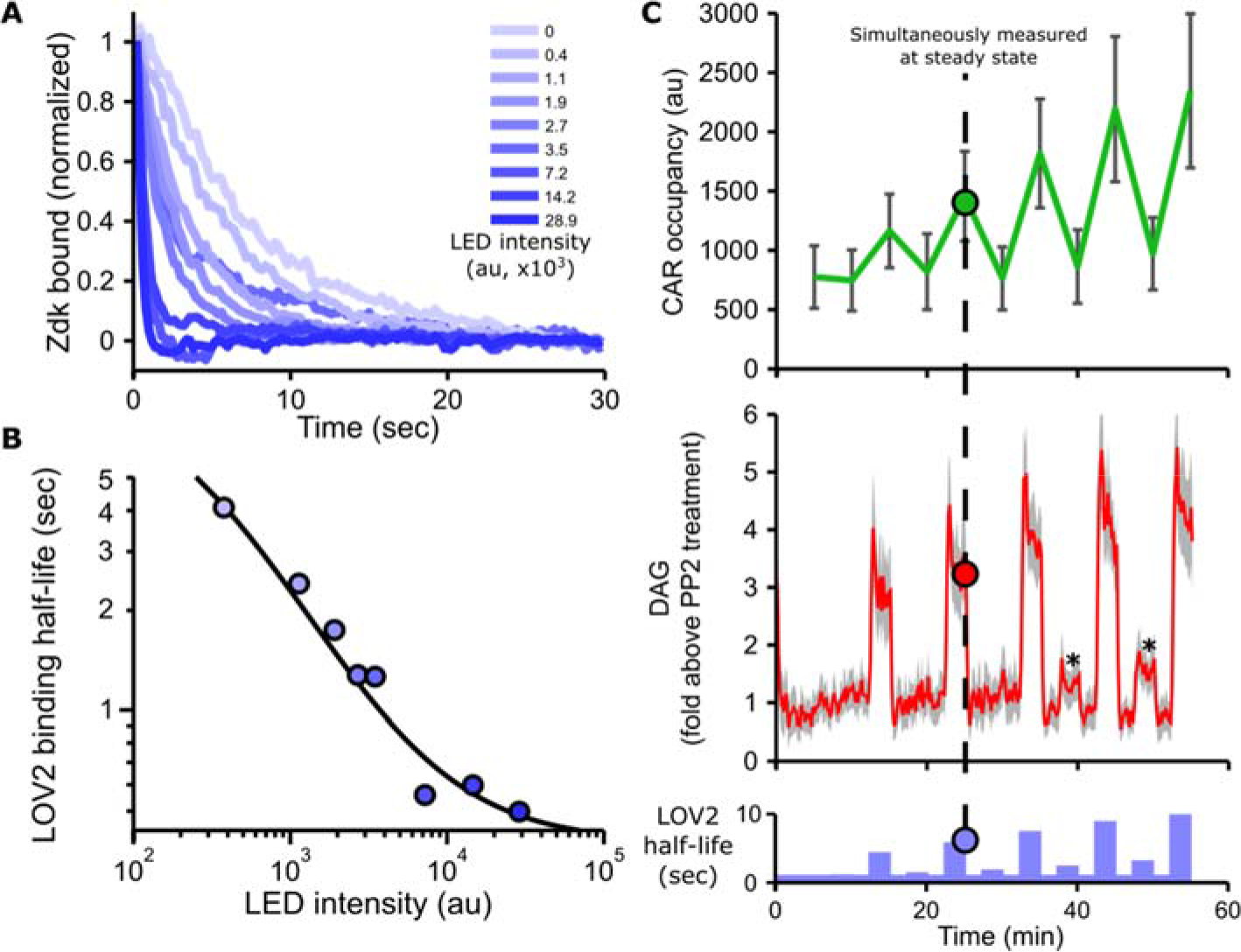
Blue light intensity titrates binding half-life, CAR occupancy and DAG levels. (**A**) *In vitro* measurements of blue light intensity-based control of LOV2-Zdk binding half-life. SLBs functionalized with LOV2 were combined with soluble, dye-labeled Zdk. After washing out free Zdk, Zdk dissociation was measured upon acute illumination with different intensities of blue light using TIRF microscopy. (**B**) Blue light intensity enforces LOV2-Zdk binding half-lives from ten seconds to hundreds of milliseconds. Binding half-lives were determined by fitting a single exponential decay to the traces shown in **A**. Data was fit using a two-step unbinding model (solid black line), consisting of light-dependent excitation of LOV2 followed by light-independent release of Zdk. (**C**) Time course showing that intermediate light levels modulate ligand binding half-lives (bottom), titrate receptor occupancy (top), and induce DAG accumulation in Jurkat cells (middle). Asterisks in middle panel highlight small but detectable increases in DAG levels to weak stimuli. n = 31 cells. Mean with 95% CI (two-sided Student’s t-test).

In addition to providing control over binding half-life, manipulating the blue light levels also finely tuned DAG levels and receptor occupancy (**Figure 2C**, **Figure 2–figure supplement 2**, **Figure 2–figure supplement 3**, Video 6 and Video 7). These data show our optogenetic input is highly titratable and directly controls ligand binding half-life.

Because increasing ligand binding half-life also increases receptor occupancy, it is difficult to separate the involvement of occupancy and half-life on downstream signaling from a single time course under one set of conditions. However, the influences of half-life and occupancy can be decoupled by keeping the sequence of blue-light stimulation the same but varying the absolute concentration of the LOV2 ligand on the SLB. One cell exposed to a low concentration of LOV2 and a long binding half-life (via low intensity blue light) can have the same receptor occupancy as another cell exposed to a high concentration of LOV2 and a short binding half-life (via higher intensity blue light; **Figure 3A**). Comparing cell signaling in this context is a very sensitive way to detect the effects of binding half-life alone, as receptor occupancy and all other ligand-intrinsic factors (including the bond’s stability under tension) are held constant.

**Figure 3.**
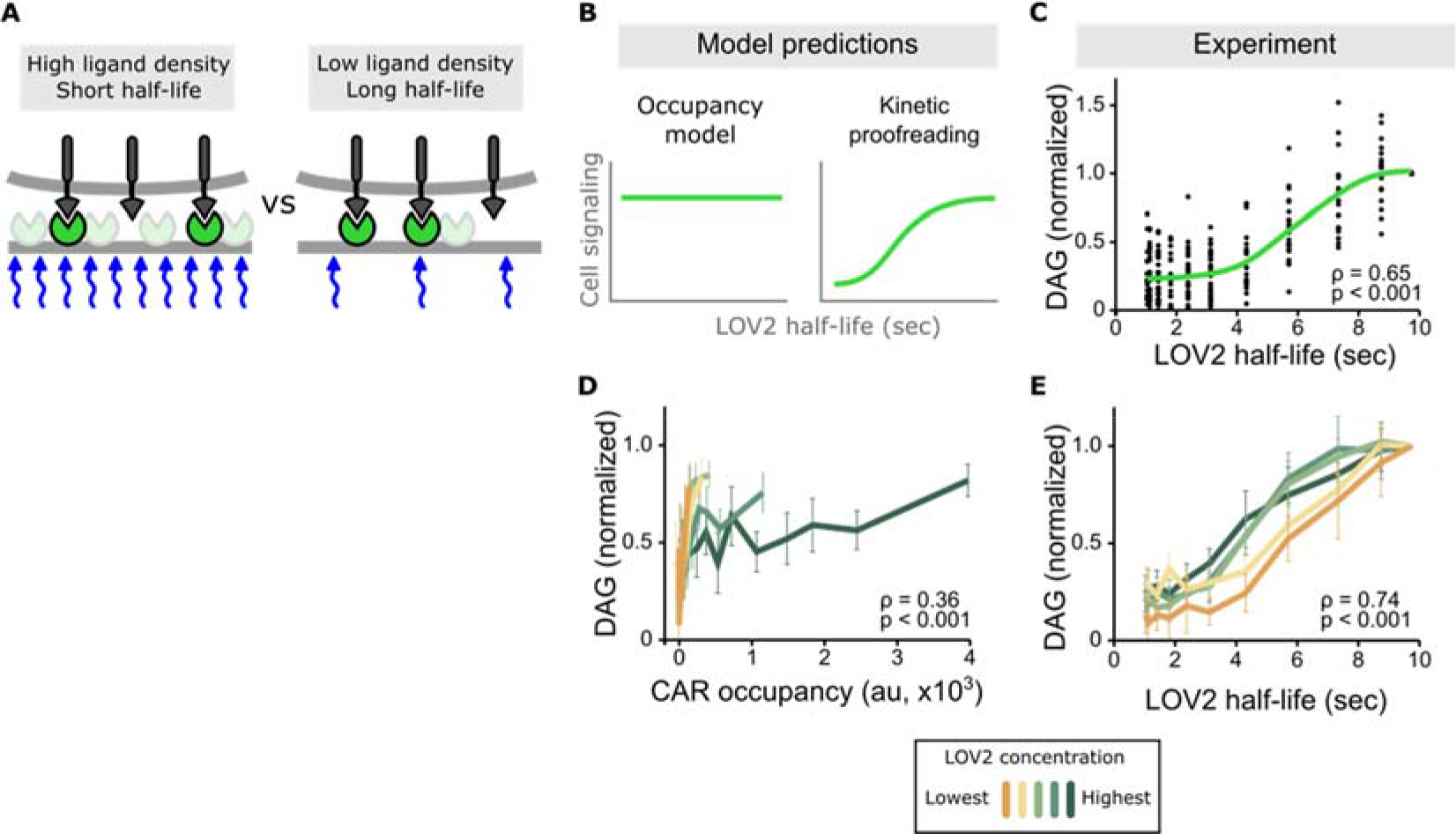
Binding half-life, not receptor occupancy, dominates CAR signaling. (**A**) A cell exposed to a high LOV2 density but a short binding half-life can have the same receptor occupancy as a cell exposed to a low LOV2 density but a long binding half-life. (**B**) At constant receptor occupancy, an occupancy model predicts binding half-life should have no effect on signaling, while kinetic proofreading predicts that increasing binding half-life should increase signaling. (**C**) At constant receptor occupancy, increasing ligand binding half-life increases DAG signaling, as shown by both non-parametric kernel smoothing regression (green line) and Spearman’s correlation coefficient (ρ). Each black dot is a single cell measurement obtained from multiple experiments over a range of LOV2 concentrations. (**D** and **E**) During individual experiments, there is a fixed concentration of LOV2, meaning that both receptor occupancy and binding half-life change in response to blue-light. We measured Spearman’s correlation coefficient to ask whether DAG levels were best described as a function of receptor occupancy or binding half-life across different LOV2 concentrations. DAG levels are more correlated with binding half-life (**E**) than they are with CAR occupancy (**D**). This is reflected in the fact that the DAG response curves nearly overlap with each other when plotted as a function of binding half-life, indicating changing the concentration of LOV2 has little effect on downstream signaling. These data are consistent with binding half-life, not receptor occupancy, dominating CAR signaling. Plotted is the mean with a 95% CI (two-sided Student’s t-test).

Conducting multiple experiments with different LOV2 concentrations and gating the data over a narrow range of receptor occupancy shows a clear result: increasing ligand binding half-life increases DAG levels, despite cells having near identical receptor occupancy (**Figure 3B, C** and **Figure 3–figure supplement 1**). Intriguingly, signaling increases the most for binding half-lives between 4-7 seconds, in close agreement with previous estimates of the binding half-life threshold for stimulatory versus non-stimulatory pMHCs (*21*–*23*).

Stimulating cells with the same intensities of blue light but different concentrations of LOV2 enabled us to measure the effects of binding half-life in the context of low, medium, and high receptor occupancy. Strikingly, the enforced ligand binding half-life predicted DAG levels well, regardless of the amount of ligand present. When DAG is plotted as a function of binding half-life, the response curves from different experiments nearly overlap with each other, showing a strong correlation coefficient (**Figure 3D, E** and **Figure 3–figure supplement 2**, top). By contrast, plotting DAG as a function of receptor occupancy shows a poor correlation (**Figure 3D** and **Figure 3–figure supplement 2**, bottom). Because half-life is a strong predictor of DAG levels across a range of receptor occupancies, these data argue that binding half-life is a major determinant of CAR signaling.

Combining the data from all experiments over a range of LOV2 ligand concentrations, we sought to quantify how strongly binding half-life influences CAR signaling. In kinetic proofreading, the delay between ligand binding and downstream signaling is modeled as a series of discrete steps (*10*). The stronger the proofreading, the greater the number of steps and the more binding half-life dominates downstream signaling. To minimize the number of assumptions, we modeled the DAG response as a saturable system downstream of “strong” proofreading steps, i.e. the forward biochemical reaction is sufficiently slow that the probability of completing a step is proportional to the ligand binding half-life. In such a model,

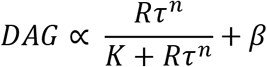

where *R* is the receptor occupancy, *τ* is the enforced ligand binding half-life, *n* is the degree of proofreading, *K* is the amount of upstream signal that generates a half-maximal DAG response, and β is basal signaling through the pathway.

The value of *n* measures how strongly ligand binding half-life affects signaling. For *n* = 0, signaling depends only on receptor occupancy and is unaffected by binding half-life. For *n* > 0, there is some degree of kinetic proofreading. As *n* increases, ligand binding half-life has a larger and larger impact on downstream signaling, to the point that short lived ligands cannot generate as much signal as long-lived ligand simply by increasing concentration and receptor occupancy (**Figure 4A**, dotted line). This aligns with what has been observed with T cells: abundant self-pMHCs do not activate T cells, while a few foreign-pMHCs can activate T cells (*4*, *24*–*27*).

**Figure 4.**
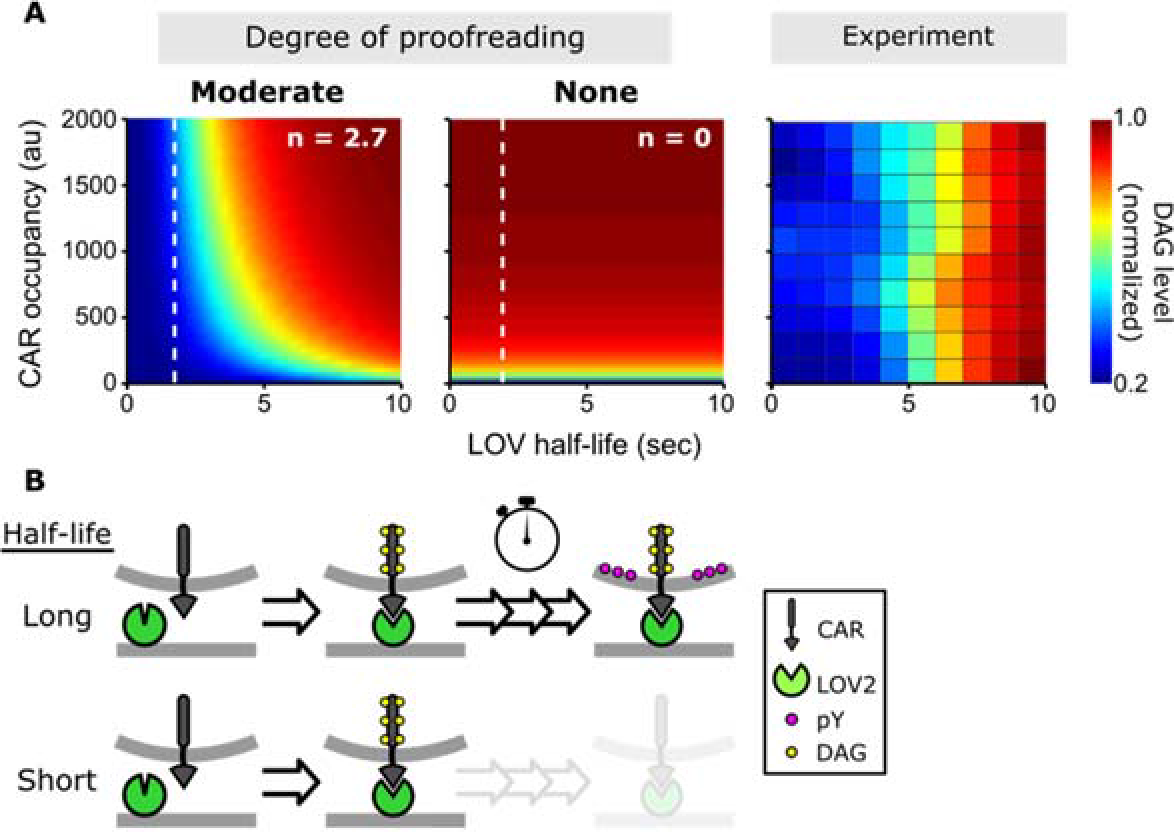
A kinetic proofreading model best explains T cell signaling. (**A**) Models for how CAR occupancy and binding half-life affect T cell signaling in the presence of moderate (left) or no kinetic proofreading (middle). To facilitate visualization, single cell measurements were fit with non-parametric kernel smoothing regression and plotted as a heat map (right). The degree of proofreading is denoted by *n*, and the value of *n* for the moderate proofreading scenario is derived from single cell measurements from all three data sets. (**Figure 4–figure supplement 1**). Experimental data of DAG levels as a function of CAR occupancy and binding half-life (right) is consistent with moderate kinetic proofreading. (**B**) Schematic of our kinetic proofreading model. After a ligand binds the receptor, it must remain bound sufficiently long to accommodate at least three slow proofreading steps. Long lived ligands survive this slow waiting period and produce strong downstream signaling, while short lived ligands dissociate and produce weak downstream signaling.

Fitting our data, we find *n* = 2.7 ± 0.5 (95% CI; **Figure 4–figure supplement 1 Figure 4–figure supplement 2)**, which means this degree of proofreading could allow T cells to amplify a 10-fold difference in ligand binding half-life into a ~5,000-fold difference in DAG levels. While this result is only true when the signaling pathway is far from saturation, it is reasonable that T cells operate far from saturation when they encounter foreign-pMHCs, given the non-stimulatory nature of self-pMHCs. The parameter *n* can also be interpreted as the minimum number of proofreading steps in the pathway, indicating there at least three steps (between CAR ligation and DAG signaling) whose forward reaction is in kinetic competition with ligand dissociation (**Figure 4B**).

How much of ligand discrimination do our results account for? As T cells can respond to between 1-10 foreign-pMHCs (*4*, *28*) in the context of ~10^5^ self-pMHCs (*1*–*3*), foreign-pMHCs must signal at least 10^4^-10^5^ fold stronger than self-pMHCs. Given that differences in binding half-lives between self-pMHCs and foreign-pMHCs are on the order of 10-fold (*7*–*9*, *14*), our observed degree proofreading can account for much, but not all, of ligand discrimination. T cells likely use kinetic proofreading in combination with other mechanisms, such as co-receptor involvement (*4*), mechanical forces (*15*, *16*, *18*), or signal amplification downstream of DAG (*29*), to fully discriminate self from non-self.

What biochemical steps underlie the observed kinetic proofreading? We addressed this question by measuring one of the earliest signaling events: the recruitment of Zap70 to the CAR. Zap70 binds to doubly phosphorylated ITAMs in the CAR's cytoplasmic domain. In contrast to DAG levels, Zap70 recruitment showed no detectable evidence of kinetic proofreading (**Figure 4–figure supplement 3)**. In other words, Zap70 recruitment was solely dependent on CAR occupancy and was unaffected by the ligand binding half-life. As live cell reporters reflect the cumulative effect of all upstream signaling steps, these results argue that the observed kinetic proofreading occurs somewhere downstream of Zap70 recruitment and upstream of DAG production.

This result is surprising, as it shows that ligand discrimination is not complete by the time the CAR signals to downstream components. Rather, proximal signaling steps downstream of ITAM phosphorylation are essential for kinetic proofreading in our system. This explanation runs counter to the conventional view that the proofreading steps correspond to the sequential phosphorylation of the ITAMs and that the TCR alone fully discriminates pMHCs, passing its decision to the proximal signaling steps. In contrast, our data argue that the proximal signaling steps kinetically discriminate the ligands while ITAM phosphorylation is insensitive to the ligand binding half-life.

While our results cannot exclude the possibility that TCR ITAM phosphorylation is sensitive to ligand binding half-life, they strongly suggest that the TCR does not fully discriminate ligands on its own. This is consistent with data showing that some Zap70 is associated with phosphorylated ITAMs in mouse thymocytes *in vivo*, even when they are not stimulated (*30*, *31*). Thus, even in primary cells, ligand discrimination is not complete at the level of Zap70 recruitment.

Even if the proofreading steps do not occur at the level of receptor activation, they still need to occur in physical proximity to the receptor so that they terminate quickly upon ligand dissociation. Candidate biochemical steps include the phosphorylation and activation of Zap70 (*32*, *33*) and/or the phosphorylation of LAT. Lck was recently shown to act as a bridge between Zap70 and LAT (*34*), suggesting that at least some fraction of LAT molecules could be tethered to phosphorylated ITAMs. Future experiments should focus on how mutations that change the kinetics of these biochemical steps affect ligand discrimination.

In summary, we addressed a fundamental question in immunology: How do T cells discriminate ligands? We overcame previous technical limitations by developing a new optogenetic approach that directly tunes ligand binding half-life while keeping constant all other parameters that could affect CAR signaling. We found direct evidence that ligand binding half-life strongly controls downstream DAG levels but not upstream signaling events, like Zap70 recruitment. Additionally, our experiments with a CAR revealed a similar degree of kinetic proofreading as complementary optogenetic experiments that signaled through the TCR (*see the co-submitted Schamel group paper*). These results argue against the common view that kinetic proofreading occurs exclusively at the level of TCR ITAM phosphorylation and instead provide evidence that it can emerge from steps in the proximal signaling pathway.

To date, optogenetics has been a powerful approach because it tells us how the timing, amount, and localization of an ensemble of signaling molecules modulate cell signaling (*35*). With the direct control over the persistence of individual protein-protein interactions presented here, optogenetics can advance to explore how the lifetime of individual molecular interactions regulates cell signaling. This could be a powerful angle for investigating other pathways where the individual molecular kinetics rather than the ensemble average are thought to gate downstream signaling (*36*–*38*).

## Acknowledgements

We thank Dyche Mullins and Peter Bieling for help with biochemical purifications, Jay Groves, Geoff O’Donoghue and Scott Hansen for help with the supported lipid bilayers, and Sam Lord for help with RICM. Thanks to Klaus Hahn, Orrin Stone, Torsten Wittmann, and Jeffrey van Haren for help troubleshooting the LOVTRAP system. Jared Toettcher, Elliot Dine, Hana El-Samad and Art Weiss provided critical reading of the manuscript.

## Author contributions

*Conceptualization*: D.T. and O.D.W.; *Methodology*: D.T.; *Investigation*: D.T.; *Resources*: O.D.W.; *Visualization*: D.T.; *Writing – Original Draft*: D.T. and O.D.W.; *Writing – Review & Editing*: O.D.W. and D.T.; Funding acquisition: O.D.W. and D.T.; *Software*: D.T.

## Data and materials availability

All data is available in the main text or the supplementary materials.

